# Inhaled nitric oxide as a rescue therapy in rat crush syndrome: translating bench research to field application

**DOI:** 10.64898/2026.03.09.710439

**Authors:** Isamu Murata, Jun Kobayashi, Shinsuke Ishihara, Nobuo Iyi

## Abstract

Crush syndrome (CS) is characterised by ischaemia/reperfusion-induced rhabdomyolysis, leading to systemic inflammation and high mortality. Building on our previous findings that intravenous nitric oxide (NO) donors improve survival in this condition, we investigated the therapeutic efficacy of inhaled NO delivered via a portable, controlled-release device in an experimental rat model of CS. Anaesthetised rats underwent bilateral hindlimb compression using rubber tourniquets for 5 h, followed by reperfusion. Among the various inhalation conditions tested, administration of NO (160 parts per million) for 2 h after reperfusion significantly increased survival rate from 20 to 90%. Improvements in haemodynamic parameters, biochemical markers, and histopathological findings correlated with enhanced survival outcomes. These results suggest that on-site NO inhalation therapy may serve as an effective first-line, emergency intervention for CS, particularly in disaster settings.

**Graphical abstract:** 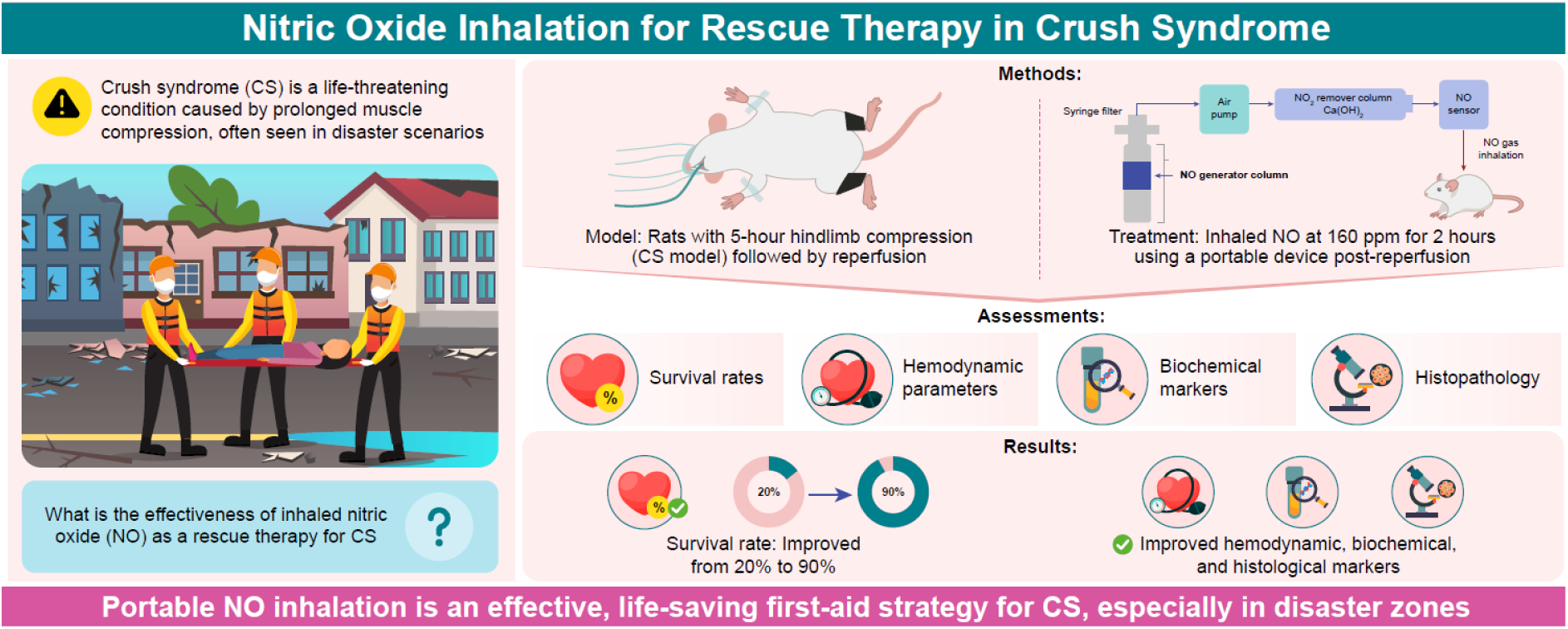

---

Crush syndrome (CS) is a potentially fatal condition that occurs following rescue from disasters, such as earthquakes^1^, landslides^2^, vehicle accidents^3^, and combat scenarios^4^. The sudden relief of prolonged and continuous compression exerted by heavy debris on lower limb skeletal muscles induces rhabdomyolysis, subsequently leading to circulatory shock, metabolic acidosis, acute respiratory distress syndrome (ARDS), and acute renal failure—complications frequently result in death^4,5^.

The aetiology of rhabdomyolysis in CS has been well characterised^5^. Compressive injury to skeletal muscle fibres imposes mechanical stress on the sarcolemma, activating non-selective stretch-activated channels. This facilitates an influx of Na^+^ and Ca^2+^ into the myocyte cytosol, triggering cellular swelling and calcium-dependent autolysis^4^. Simultaneously, local ischaemia induces a metabolic shift from aerobic to anaerobic pathways, resulting in ATP depletion and subsequent muscle cell damage^6^, culminating in rhabdomyolysis.

Importantly, rhabdomyolysis does not develop during ischaemia per se but rather upon reperfusion of the injured muscle^4^. During reperfusion, oxidative stress leads to muscle cell lysis and the release of intracellular constituents—such as potassium and myoglobin—into the circulation, precipitating early and late fatal complications, including cardiac arrhythmias, myoglobinuric renal failure, and systemic inflammation^5^.

Various animal models of CS have been established to simulate skeletal muscle compression–decompression injury^7,8^, and the therapeutic effects of multiple pharmacological agents have been investigated^9–11^. We have previously developed a simple rat model using bilateral hindlimb compression with rubber tourniquets to study CS pathophysiology and explore treatment strategies^5^. Using this model, we demonstrated the survival benefit of intravenous nitrite—a nitric oxide (NO) donor— highlighting its anti-inflammatory properties in CS treatment^12^.

In the present study, we evaluated the potential of NO inhalation as an alternative therapeutic modality^13^. Following ischaemia–reperfusion (I/R) injury, lysates derived from damaged skeletal muscle accumulate in the pulmonary vasculature, triggering inflammation and respiratory failure^5^. The lungs—being the first capillary bed to receive systemic circulation—serve as a primary site of injury, often culminating in ARDS. Inhaled NO has been shown to attenuate pulmonary endothelial inflammation and limit the downstream systemic propagation of inflammatory mediators^14^. Moreover, recent studies indicate that inhaled NO confers protection against myocardial I/R injury by forming stable *S*-nitrosothiols (RSNOs) in plasma and erythrocytes during pulmonary circulation^15–17^. These RSNOs may exert NO-dependent signalling effects in extrapulmonary organs, including the kidney and injured skeletal muscles.

Here, we employed a portable, controlled-release NO gas delivery device developed by Ishihara and Iyi—co-authors of the present study. They have previously described its technical specifications and proposed its clinical utility in emergency scenarios^18^. The findings of this study highlight its substantial potential for use in first-aid and emergency interventions, particularly in field settings.

## Results

### Effects of ischaemic duration, inhaled NO concentration, and timing of NO inhalation on survival in a rat model of CS

In our previous study, we established that 5 h of bilateral hindlimb ischaemia induced by rubber tourniquets produced the highest mortality in a rat model of CS; therefore, this duration was selected for the present experiments^5^. To determine the optimal concentration of inhaled NO for therapeutic intervention, we compared the effects of low (20 parts per million [ppm]) and high (160 ppm) concentrations of inhaled NO on survival outcomes in the CS rat model. We then evaluated the impact of the timing of 20 ppm NO inhalation using various regimens relative to reperfusion: 1 h or 2 h before reperfusion, 1 h before and 1 h after reperfusion, and 1 h or 2 h after reperfusion. Untreated CS rats exhibited high mortality, with survival rates of approximately 20% at both 24 and 48 h after reperfusion. Notably, rats receiving 2 h of inhaled NO beginning after reperfusion demonstrated an improved survival rate of approximately 40% at 48 h. In contrast, all other NO inhalation regimens yielded survival rates comparable to those of the untreated group, remaining around 20% (Fig. 1a).

**Fig. 1.**
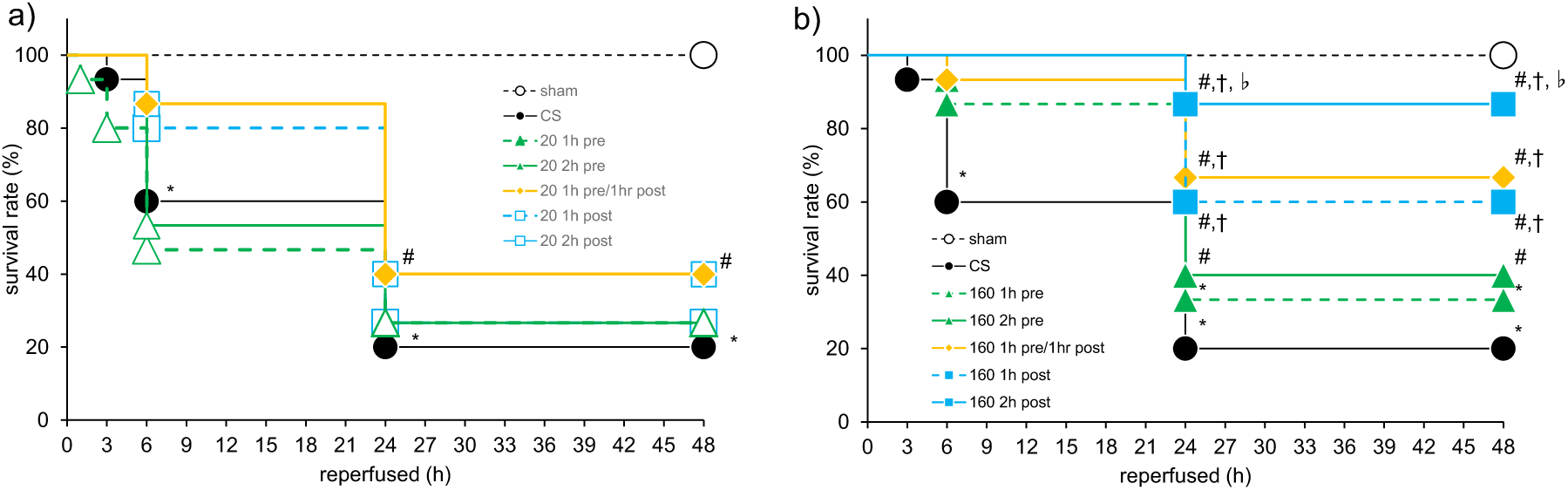
Survival rates with nitric oxide inhalation. a) 20 ppm nitric oxide inhalation. Survival rates of sham-operated rats, CS rats, and CS rats inhaling 20 ppm NO (20 1 h pre, 20 2 h pre, 20 1 h post, 20 2 h post, and 20 1 h pre/1 h post) were assessed at 0, 1, 3, 6, 24, and 48 h after reperfusion. Survival curves are generated using the Kaplan–Meier method (n=15 per group). \**p*<0.05 vs sham group, #*p*<0.05 vs CS group (log-rank test). b) 160 ppm nitric oxide inhalation. Survival rates of sham-operated rats, CS rats, and CS rats inhaling 160 ppm NO (160 1 h pre, 160 2 h pre, 160 1 h post, 160 2 h post, and 160 1 h pre/1 h post) at 0, 1, 3, 6, 24, and 48 h after reperfusion. *Abbreviations:* NO, nitric oxide; CS, crush syndrome; ppm, parts per million; 20 1 h pre, 20 ppm NO inhalation 1 h before reperfusion; 20 2 h pre, 20 ppm NO inhalation 2 h before reperfusion; 20 1 h post, 20 ppm NO inhalation 1 h after reperfusion; 20 2 h post, 20 ppm NO inhalation 2 h after reperfusion; 20 1 h pre/1 h post, 20 ppm NO inhalation 1 h before and 1 h after reperfusion. 160 1 h pre, 160 ppm NO inhalation 1 h before reperfusion; 160 2 h pre, 160 ppm NO inhalation 2 h before reperfusion; 160 1 h post, 160 ppm NO inhalation 1 h after reperfusion; 160 2 h post, 160 ppm NO inhalation 2 h ter reperfusion; 160 1 h pre/1 h post, 160 ppm NO inhalation 1 h before and 1 h after reperfusion.

Compared with the survival rates observed in the 20 ppm NO inhalation groups, inhalation of 160 ppm NO—particularly when administered after reperfusion—markedly improved survival outcomes. Specifically, survival rates were 60%, 70%, and 90% for NO inhalation administered for 1 h after reperfusion, 1 h before plus 1 h after reperfusion, and 2 h after reperfusion, respectively. These rates were significantly higher than those observed in the 160 ppm NO groups administered 1 h or 2 h before reperfusion (30% and 40%, respectively) and in the CS control group (20%). Among all experimental groups, 160 ppm NO inhalation for 2 h after reperfusion yielded the highest survival rate (90%), second only to the sham group, which showed 100% survival (Fig. 1b). These findings indicate that higher concentrations of inhaled NO (160 ppm vs 20 ppm), longer durations of administration (2 h vs 1 h), and after reperfusion delivery rather than before reperfusion are key determinants of improved survival in this CS rat model. Accordingly, subsequent experiments were conducted using 160 ppm NO inhalation to evaluate its therapeutic effects on haemodynamic parameters, biochemical markers, and histopathological changes.

### Effects of NO inhalation on electrocardiogram parameters in CS rats

Supplementary Figs. S1a–e show electrocardiogram (ECG) parameters from sham-operated rats, CS controls, and NO-inhaled CS rats (2 h before, 1 h before /1 h after, and 2 h after reperfusion) assessed at 3, 24, and 48 h after reperfusion. Compared with the sham group, the QRS, PR, and QT intervals were significantly prolonged in CS control rats at all time points (Supplementary Figs. S1a–c). In contrast, NO inhalation attenuated these interval prolongations, with the most pronounced improvements observed in rats receiving NO for 2 h after reperfusion (Supplementary Figs. S1a–c). In CS control rats, tall and peaked T wave amplitudes—so-called tented T waves indicative of hyperkalaemia—were evident at 3, 24, and 48 h after reperfusion (Supplementary Fig. S1e). NO inhalation significantly reduced T-wave amplitudes, again most prominently in the 2 h after reperfusion group. Additionally, although P-wave amplitude was reduced in CS control rats compared with sham rats, it was restored by NO inhalation, with the greatest recovery observed in the 2 h after reperfusion group (Supplementary Fig. S1d). Collectively, these results indicate that NO inhalation mitigates I/R-induced electrophysiological abnormalities, including hyperkalaemia-associated T-wave changes (Supplementary Fig. S1e), atrioventricular nodal conduction delays (Supplementary Fig. S1b), and atrial myocardial dysfunction (Supplementary Figs. S1a, c, d).

### Effects of NO inhalation on haemodynamic and biochemical parameters in CS rats

In CS rats, heart rate (HR) and blood pressure (BP) parameters—including systolic blood pressure (SBP), diastolic blood pressure (DBP), and mean blood pressure (MBP)—were significantly lower than those in the sham group. In contrast, NO inhalation significantly improved BP in all treated groups (2 h before, 1 h before /1 h after, and 2 h after reperfusion). Restoration of BP to near-sham levels was most pronounced in rats receiving NO for 2 h after reperfusion (Table 1).

**Table 1.**
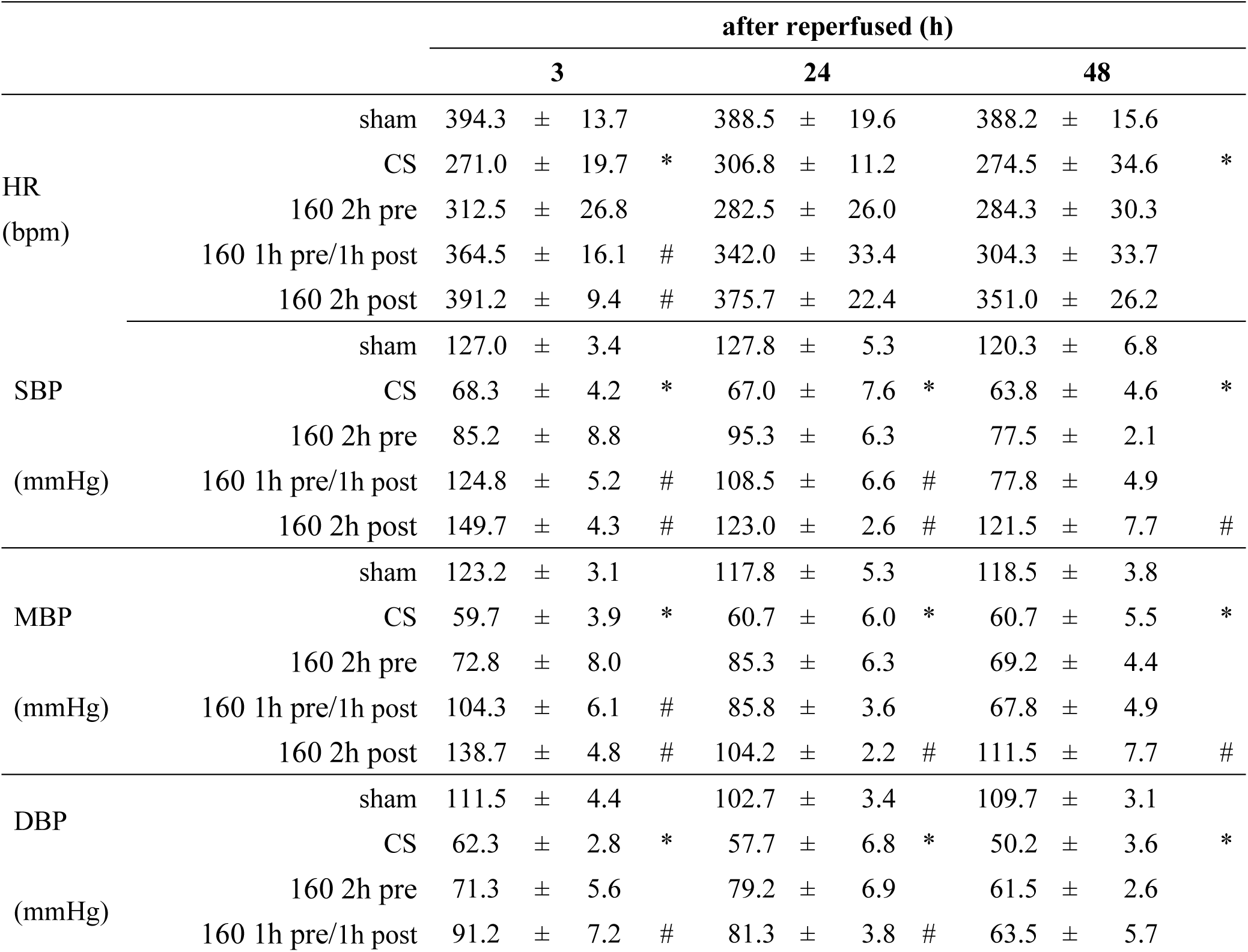
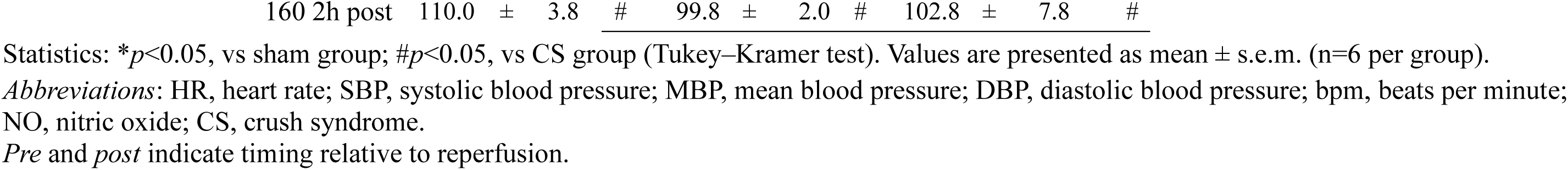
Effects of 160 ppm NO inhalation on heart rate and blood pressure in a rat model of crush syndrome.

Table 2 summarises the time course of plasma biochemical parameters in sham-operated rats, CS control rats, and NO-treated CS rats. Levels of creatine kinase (CK), myoglobin (Mb), glucose (Glu), potassium (K^+^), blood urea nitrogen (BUN), and creatinine (Cre) were significantly elevated in CS control rats following skeletal muscle I/R injury, consistent with rhabdomyolysis, hyperkalaemia, and acute renal dysfunction. NO inhalation significantly attenuated these elevations, with the greatest reductions observed in the 2 h after reperfusion group. Collectively, these findings indicate that after reperfusion NO inhalation provides superior protection against I/R-induced rhabdomyolysis, electrolyte imbalance, and renal impairment.

**Table 2.** Effects of 160 ppm NO inhalation on biochemical parameters in a rat model of crush syndrome.

|  |  | after reperfused (h) |  |  |  |  |  |  |  |  |  |  |  |
| --- | --- | --- | --- | --- | --- | --- | --- | --- | --- | --- | --- | --- | --- |
|  |  | 3 |  |  |  | 24 |  |  |  | 48 |  |  |  |
| CPK<br>(×10<sup>4</sup> IU/L) | sham | 0.2 | ± | 0.0 |  | 0.2 | ± | 0.0 |  | 0.2 | ± | 0.0 |  |
|  | CS | 6.8 | ± | 1.4 | * | 20.6 | ± | 3.2 | * | 22.2 | ± | 1.8 | * |
|  | 160 2h pre | 4.4 | ± | 0.9 |  | 18.2 | ± | 0.8 |  | 16.9 | ± | 2.6 |  |
|  | 160 1h pre/1h post | 3.4 | ± | 0.7 | # | 12.0 | ± | 1.6 | # | 12.1 | ± | 1.4 | # |
|  | 160 2h post | 3.0 | ± | 0.3 | # | 8.0 | ± | 0.6 | # | 6.6 | ± | 0.6 | # |
| Myoglobin<br>(mg/mL) | sham | 10.8 | ± | 2.2 |  | 4.8 | ± | 0.2 |  | 4.7 | ± | 0.2 |  |
|  | CS | 246.5 | ± | 56.6 | * | 512.2 | ± | 58.1 | * | 566.0 | ± | 82.4 | * |
|  | 160 2h pre | 238.3 | ± | 53.5 |  | 446.0 | ± | 22.8 |  | 504.0 | ± | 38.2 |  |
|  | 160 1h pre/1h post | 227.7 | ± | 34.2 |  | 318.3 | ± | 36.7 |  | 332.3 | ± | 14.4 |  |
|  | 160 2h post | 150.2 | ± | 15.0 | # | 302.5 | ± | 25.1 | # | 306.0 | ± | 39.1 | # |
| Glu<br>(mg/dL) | sham | 155.5 | ± | 6.9 |  | 170.2 | ± | 9.1 |  | 181.0 | ± | 6.1 |  |
|  | CS | 224.7 | ± | 21.8 | * | 257.3 | ± | 22.4 | * | 260.2 | ± | 24.1 | * |
|  | 160 2h pre | 195.2 | ± | 21.2 |  | 211.2 | ± | 7.8 |  | 233.8 | ± | 13.7 |  |
|  | 160 1h pre/1h post | 183.2 | ± | 10.1 |  | 204.2 | ± | 29.3 |  | 214.0 | ± | 29.3 |  |
|  | 160 2h post | 176.7 | ± | 18.2 |  | 190.3 | ± | 26.2 |  | 242.8 | ± | 36.5 |  |
| pottasium<br>(mEq/L) | sham | 3.93 | ± | 0.13 |  | 4.47 | ± | 0.08 |  | 4.03 | ± | 0.14 |  |
|  | CS | 6.07 | ± | 0.17 | * | 8.10 | ± | 0.32 | * | 6.63 | ± | 0.31 | * |
|  | 160 2h pre | 5.97 | ± | 0.15 |  | 5.93 | ± | 0.31 |  | 5.93 | ± | 0.20 |  |
|  | 160 1h pre/1h post | 5.17 | ± | 0.23 |  | 5.78 | ± | 0.31 | # | 4.82 | ± | 0.16 | # |
|  | 160 2h post | 4.77 | ± | 0.16 | # | 5.00 | ± | 0.12 | # | 3.95 | ± | 0.11 | # |
| BUN<br>(mg/dL) | sham | 20.2 | ± | 0.8 |  | 23.3 | ± | 1.7 |  | 22.5 | ± | 1.5 |  |
|  | CS | 41.2 | ± | 3.3 | * | 113.8 | ± | 6.4 | * | 113.5 | ± | 6.4 | * |
|  | 160 2h pre | 38.5 | ± | 2.6 |  | 59.0 | ± | 1.9 |  | 59.2 | ± | 4.4 |  |
|  | 160 1h pre/1h post | 24.7 | ± | 1.5 | # | 41.2 | ± | 3.0 | # | 46.5 | ± | 5.5 | # |
|  | 160 2h post | 19.0 | ± | 1.1 | # | 45.8 | ± | 8.4 | # | 30.2 | ± | 1.2 | # |
| Cre<br>(mg/dL) | sham | 0.21 | ± | 0.02 |  | 0.21 | ± | 0.02 |  | 0.22 | ± | 0.01 |  |
|  | CS | 1.59 | ± | 0.16 | * | 1.22 | ± | 0.26 | * | 1.15 | ± | 0.18 | * |
|  | 160 2h pre | 1.07 | ± | 0.21 |  | 0.95 | ± | 0.14 |  | 1.19 | ± | 0.09 |  |
|  | 160 1h pre/1h post | 0.98 | ± | 0.15 |  | 0.64 | ± | 0.20 |  | 1.10 | ± | 0.10 |  |
|  | 160 2h post | 1.13 | ± | 0.22 |  | 0.84 | ± | 0.15 |  | 0.47 | ± | 0.04 | # |

Statistics: \* $p < 0.05$ , vs sham group; # $p < 0.05$ , vs CS group (Tukey–Kramer test). Values are presented as the mean ± s.e.m. (n=6 per group).
Abbreviations: CPK, creatine phosphokinase; Glu, glucose; BUN, blood urea nitrogen; Cre, creatinine; NO, nitric oxide; CS, crush syndrome; ppm, parts per million; s.e.m., standard error of mean; *Pre* and *post* indicate timing relative to reperfusion.

### Arterial blood gas and methaemoglobin analysis in CS rats

Arterial blood gas analysis revealed persistent acidosis in CS rats at 24 and 48 h after reperfusion. NO inhalation corrected arterial pH from 7.28 in CS control rats to 7.39 in the 2 h before and 1 h before /1 h after reperfusion groups and to 7.47 in the 2 h after reperfusion group at 24 h after reperfusion. This improvement in arterial pH was sustained through 48 h (Table 3). Elevations in the anion gap and arterial lactate levels in CS rats indicated a primary metabolic acidosis, further supported by reductions in base excess (BE) and bicarbonate (HCO_3_^-^). However, the presence of hypercapnia exceeding the expected physiological respiratory compensation at both 24 and 48 h after reperfusion suggests the coexistence of mixed metabolic and respiratory acidosis.

**Table 3.**
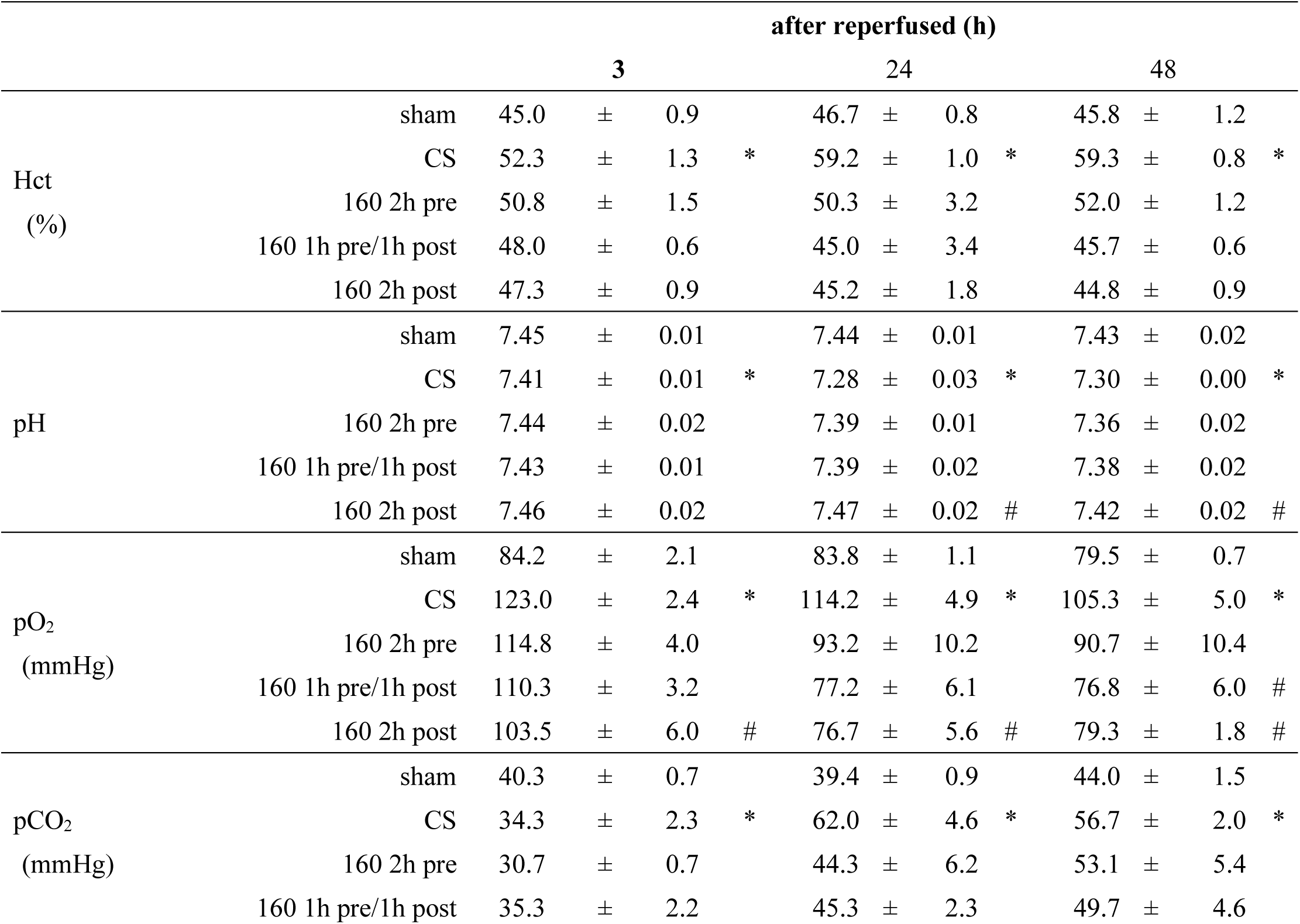
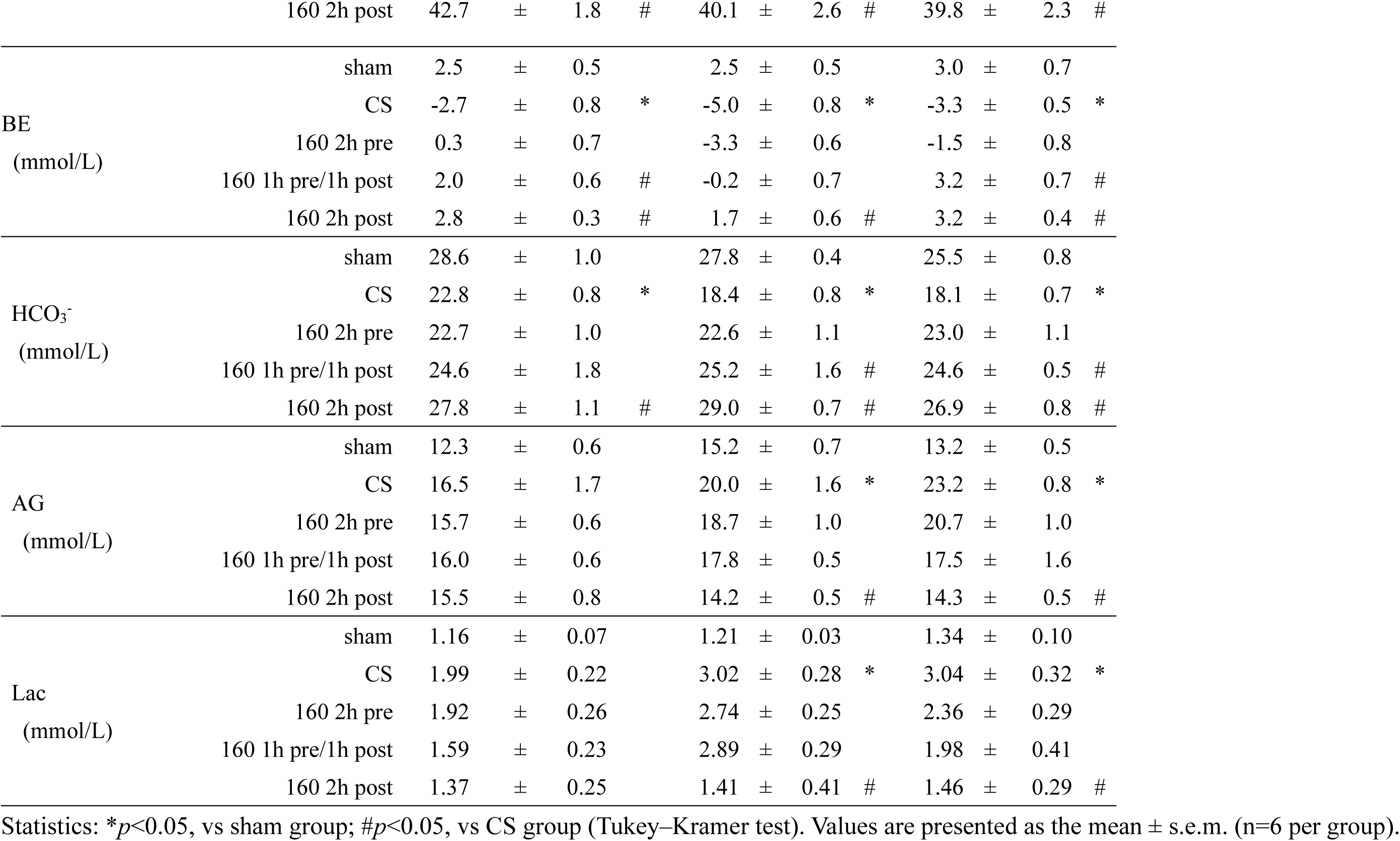
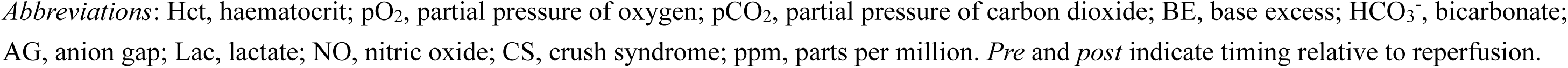
Effects of 160 ppm NO inhalation on arterial blood gas parameters in a rat model of crush syndrome.

Methaemoglobin (Met-Hb) levels were evaluated from 2 h before reperfusion through 48 h after reperfusion in sham-operated rats, CS control rats, and NO-treated CS rats (160 ppm). Met-Hb levels peaked at approximately 4% in the 2 h before and 1 h before /1 h after reperfusion groups. In contrast, rats receiving NO for 2 h after reperfusion exhibited lower Met-Hb levels, remaining below 3% throughout the observation period (Supplementary Fig. S2). These findings indicate that after reperfusion NO inhalation effectively improves acid–base balance while maintaining a relatively lower risk of NO-induced methaemoglobinaemia.

### Effects of NO inhalation on systemic cytokine profiles in CS rats

Plasma concentrations of the pro-inflammatory cytokines tumour necrosis factor-α (TNF-α), interleukin (IL)-1β, and IL-6, as well as the anti-inflammatory cytokine IL-10, were measured at 24 and 48 h after reperfusion in sham-operated rats, CS control rats, and NO-treated CS rats (160 ppm for 2 h before, 1 h before /1 h after, and 2 h after reperfusion). Compared with sham rats, CS rats exhibited significantly elevated plasma levels of TNF-α, IL-1β, and IL-6 at both time points. NO inhalation significantly attenuated these elevations, with the most pronounced reductions observed in the 2 h after reperfusion group (Fig. 2a). Although cytokine levels remained elevated at 48 h after reperfusion, peak concentrations were generally lower than those observed at 24 h, suggesting a gradual resolution of the inflammatory response. In contrast to the pro-inflammatory cytokines, IL-10 levels were similarly elevated in CS control and NO-treated groups at 24 h after reperfusion (Fig. 2a). This persistent elevation of IL-10 may reflect an endogenous anti-inflammatory response aimed at limiting I/R injury, potentially through suppression of neutrophil recruitment and cytokine production, consistent with previous reports on the regulatory role of IL-10 in myocardial and pulmonary I/Rinjury^19–21^. At 48 h after reperfusion, IL-10 levels showed a declining trend in NO-treated rats, with a more pronounced reduction in the 2 h after reperfusion group, potentially indicating progression towards resolution of the inflammatory phase.

**Fig. 2.**
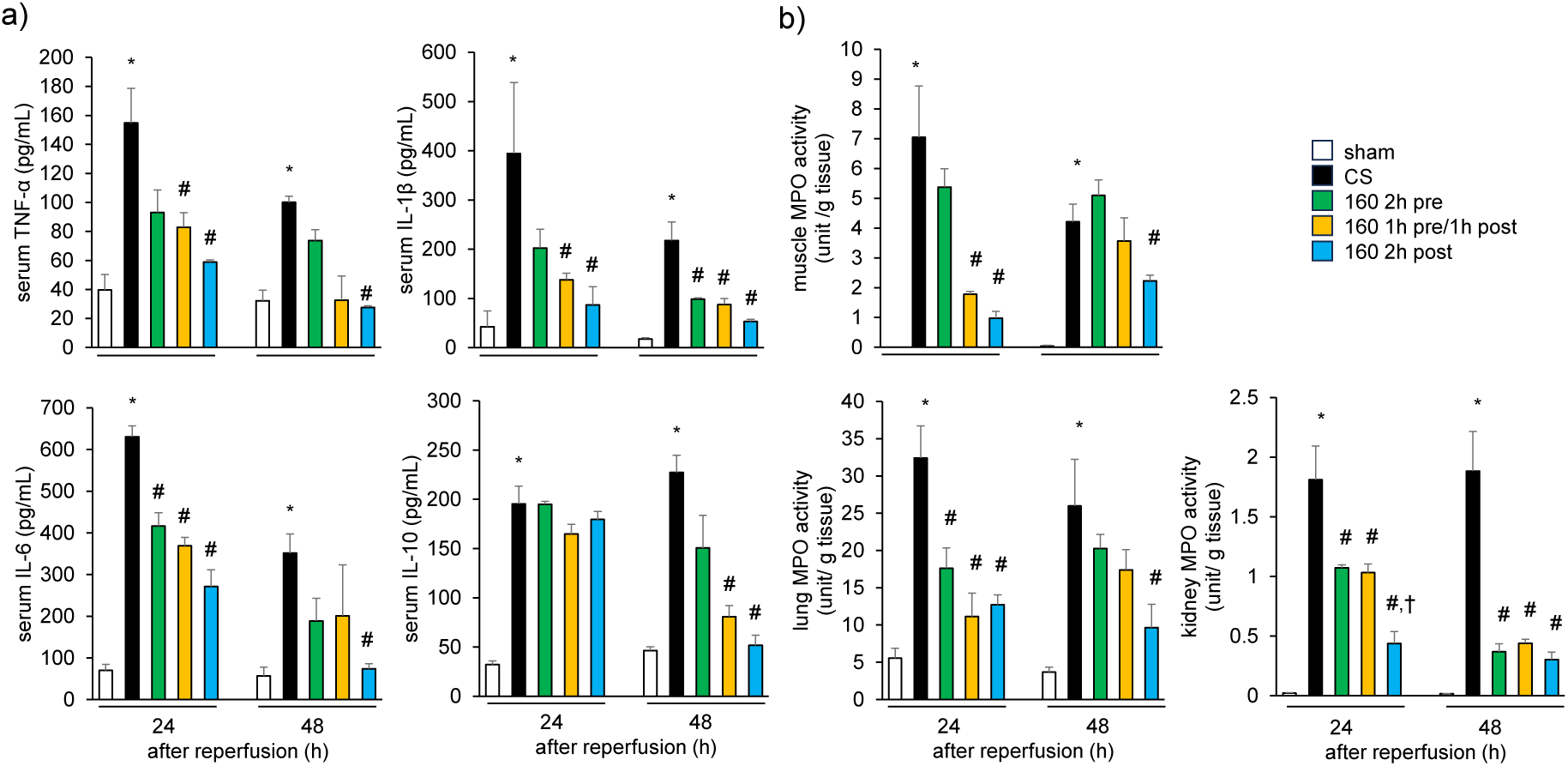
Plasma levels of inflammatory, anti-inflammatory cytokines and tissue myeloperoxidase activity levels. a) Plasma levels of inflammatory, and anti-inflammatory cytokines. Plasma levels of the pro-inflammatory cytokines, TNF-α, IL-1β, and IL-6, and the anti-inflammatory cytokine (IL-10), are measured at 24 and 48 h after reperfusion in sham-operated rats, CS control rats, and CS rats treated with inhaled NO (160 ppm). \**p*<0.05, vs sham group, #*p*<0.05, vs CS group (Tukey–Kramer test). Values are presented as the mean ± s.e.m. (n=6 per group). b) Tissue myeloperoxidase activity levels. MPO activity in skeletal muscle, lung, and kidney tissues is measured at 24 and 48 h after reperfusion in sham-operated rats, CS control rats, and CS rats treated with inhaled NO (160 ppm). \**p*<0.05, vs sham group; #*p*<0.05, vs CS group; †*p*<0.05, vs 160 ppm before reperfusion group (Tukey–Kramer test). Values are presented as mean ± s.e.m. (n=6 per group). *Abbreviations:* NO, nitric oxide; CS, crush syndrome; ppm, parts per million; TNF-α, tumour necrosis factor-α; IL-1β, interleukin-1β; IL-6, interleukin-6; IL-10, interleukin-10; s.e.m., standard error of mean. *Pre* and *post* indicate timing relative to reperfusion

### Effects of NO inhalation on tissue myeloperoxidase activity in CS rats

Myeloperoxidase (MPO) activity, a marker of leukocyte infiltration, was assessed in skeletal muscle, lung, and kidney tissues at 24 and 48 h after reperfusion in sham-operated rats, CS control rats, and NO-treated CS rats (160 ppm for 2 h before, 1 h before /1 h after, and 2 h after reperfusion). MPO activity was significantly elevated in all examined organs of CS rats compared with sham animals, reflecting enhanced leukocyte accumulation secondary to I/R injury. Inhaled NO markedly suppressed MPO activity in each organ at both time points, with the most pronounced reductions observed in the 2 h after reperfusion group (Fig. 2b). These findings are consistent with the temporal profile of increased plasma pro-inflammatory cytokines (Fig. 2a) and suggest that NO inhalation mitigates systemic and local inflammatory responses by limiting leukocyte infiltration not only in tissues directly affected by I/R injury but also in remote organs, including the lungs and kidneys. The mechanisms underlying this multi-organ anti-inflammatory effect of inhaled NO are further addressed in the Discussion section.

### Effects of NO inhalation on histopathology of the lungs, skeletal muscles, and kidneys in CS rats

In sham-operated rats, gastrocnemius muscles displayed normal histological architecture. In contrast, CS rats exhibited marked interstitial oedema, leukocyte infiltration, muscle fibre degeneration, and atrophy. These pathological changes were attenuated by NO inhalation, with the most pronounced improvements observed in rats treated for 2 h after reperfusion at both 24 and 48 h after reperfusion (Fig. 3a and b, upper panels).

**Fig. 3.**
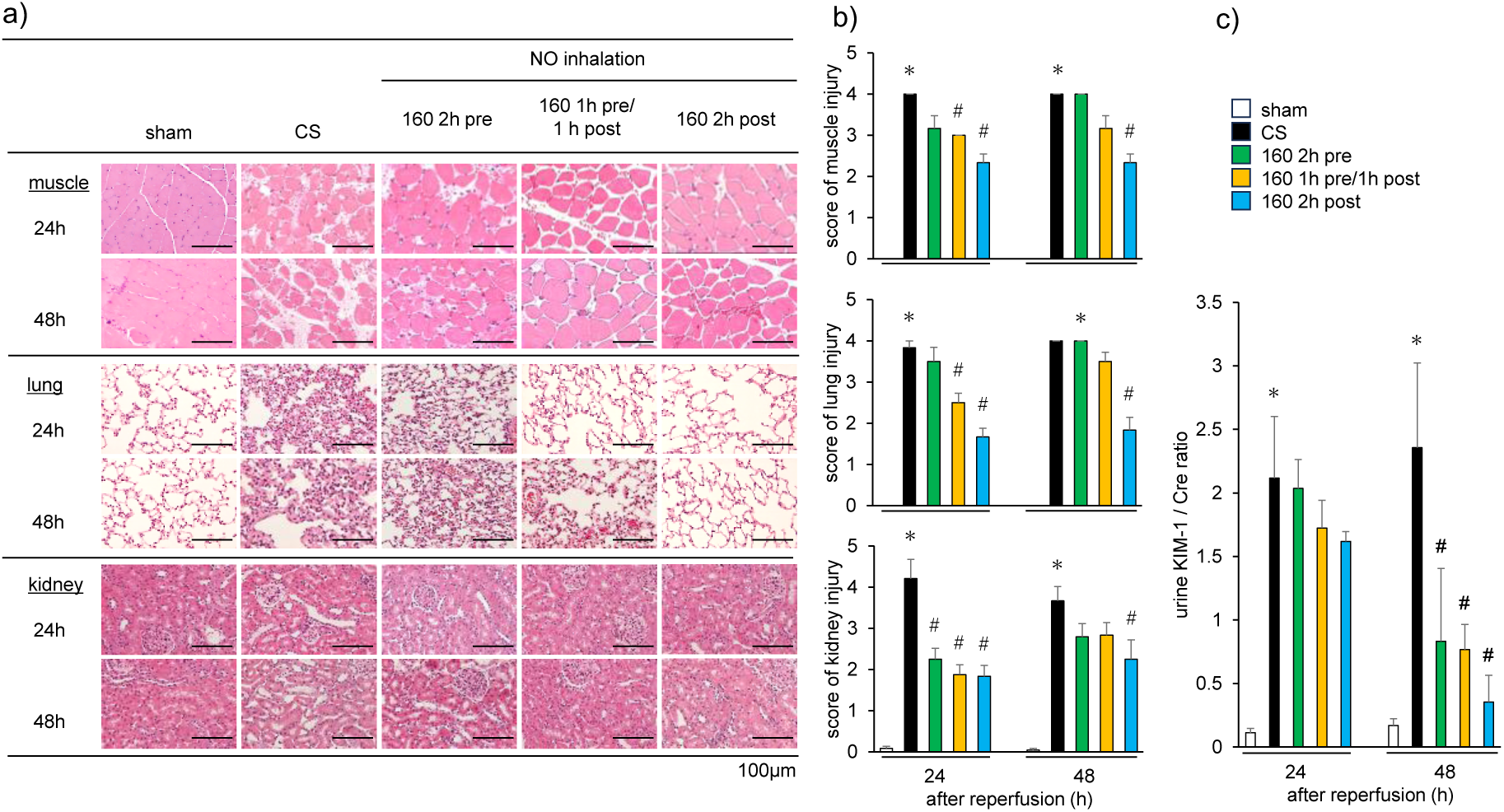
Effects of NO inhalation on the histopathology of skeletal muscle, lung, and kidney in CS rats. a) Representative histological sections of gastrocnemius muscle, lung, and kidney tissues obtained at 24 and 48 h after reperfusion from sham-operated rats, CS rats, and CS rats treated with inhaled NO (160 ppm) for 2 h before, 1 h pre/1 h post, or 2 h after reperfusion. **Upper panels:** Sham rats show normal gastrocnemius muscle architecture at both time points. CS rats exhibit marked oedema, muscle fibre degeneration, and atrophy. These pathological changes are attenuated by NO inhalation, most prominently in the 2 h after reperfusion group. **Middle panels**: Sham rats display normal alveolar architecture with thin septa. CS rats show pronounced intra-alveolar and interstitial oedema with inflammatory cell infiltration, which is reduced by NO inhalation at both 24 and 48 h after reperfusion. **Lower panels**: Sham kidneys exhibit preserved cortical architecture. CS rats show dilation of distal tubules with epithelial flattening, particularly at 48 h after reperfusion. NO inhalation mitigates these renal pathological changes at both time points. Haematoxylin and eosin staining; original magnification, ×200. c) Quantitative analysis of the variables scored in skeletal muscles, lungs and kidneys before and after NO inhalation. Quantitative analysis of histopathological variables in skeletal muscle, lung, and kidney tissues at 24 and 48 h after reperfusion in sham-operated rats, CS rats, and CS rats treated with inhaled NO (160 ppm). NO inhalation significantly improves histopathological scores across all organs, with the most pronounced therapeutic effects observed in the 2 h after reperfusion group at both time points. d) Urinary excretion of kidney injury molecule-1. Urinary levels of KIM-1 measured at 24 and 48 h after reperfusion in sham-operated rats, CS rats, and CS rats treated with inhaled NO (160 ppm). CS rats show significantly increased urinary KIM-1 excretion compared with sham rats. NO inhalation reduces KIM-1 levels, with a decreasing trend at 24 h and a significant reduction at 48 h after reperfusion, particularly in the 2 h after reperfusion group. \**p*<0.05, vs sham group; #*p*<0.05, vs CS group (Tukey–Kramer test). Values are presented as the mean ± s.e.m. (n=6 per group). *Abbreviations*: NO, nitric oxide; CS, crush syndrome; KIM-1, kidney injury molecule-1; ppm, parts per million. *Pre* and *post* indicate timing relative to reperfusion.

Similarly, histological examination of lung tissue revealed thin alveolar septa in sham rats, whereas CS rats showed extensive intra-alveolar and interstitial oedema accompanied by leukocyte infiltration. These inflammatory features were substantially reduced by NO inhalation, particularly in the 2 h after reperfusion group (Fig. 3a and 3b, middle panels).

Renal histology in sham rats showed intact glomeruli and normal proximal and distal tubular structures. In contrast, CS rats exhibited moderate dilation of distal tubules and epithelial flattening, with these lesions more prominent at 48 than at 24 h after reperfusion. NO inhalation ameliorated these pathological changes at both time points, again with greater efficacy in the 2 h after reperfusion group (Fig. 3a and b, lower panels, 3b). Consistent with these histological findings, urinary levels of kidney injury molecule-1 (KIM-1), a transmembrane protein with immunoglobulin and mucin domains and a recognised urinary marker of acute proximal tubular injury^22^, were significantly elevated in CS rats. NO inhalation reduced KIM-1 excretion, with a modest decrease at 24 h and a significant reduction at 48 h after reperfusion, particularly in rats treated with NO for 2 h after reperfusion (Fig. 3c). Quantitative scoring of histopathological variables in skeletal muscle, lung and kidney tissues corroborated these observations, demonstrating significant improvements following NO inhalation. The therapeutic effect was most pronounced in the 2 h after reperfusion group at both 24 and 48 h across all organs examined (Fig. 3a and b).

## Discussion

To the best of our knowledge, this is the first report to establish NO inhalation as a practical and effective treatment strategy for CS that bypasses the need for intravenous drug administration or fluid infusion.

CS remains a major cause of mortality at disaster sites, necessitating the urgent development of effective, field-deployable therapeutic strategies. Among emerging interventions^3^, NO donors have shown promise as protective agents against I/R injury, a central pathological component of CS. Our group has previously demonstrated that intravenous administration of nitrite, an NO donor, immediately before reperfusion significantly improves survival in a rat model of CS, increasing survival rates from 24% in untreated CS controls to 36% and 64% with 100 and 200–500 μmol/kg of NaNO_2_, respectively^12^. However, systemic hypotension—a well-recognised adverse effect of intravenous nitrite—raises safety concerns, particularly in the setting of CS-associated hypovolaemic shock. In our earlier study, intravenous nitrite administration did not restore mean arterial pressure to levels observed in sham-operated animals^12^.

To address this limitation and evaluate the translational potential of NO-based therapy in real-world disaster settings, we examined the effects of inhaled NO in a rat model of CS. We utilised a ready-to-use NO gas-releasing device, recently described by Ishihara and Iyi^18^, which offers a practical and rapidly deployable method for delivering inhaled NO in austere environments. In this study, we focused on optimising three key experimental variables—ischaemic duration, inhaled NO concentration, and timing of NO administration (2 h before, 1 h before /1 h after, and 2 h after reperfusion)—to establish conditions that closely mimic the mortality associated with CS. Accurate determination of the ischaemic period is critical for the development of effective therapeutic strategies. Belkin et al.^23^, using spectrophotometric analysis of skeletal muscle injury in a rat hindlimb tourniquet model, demonstrated that significant tissue damage begins at 3 h of ischaemia and progressively worsens with longer durations (4, 5, and 6 h). Consistently, Murata et al.^5^ reported that tourniquet-induced 5 h of bilateral hindlimb ischaemia in rats resulted in near-complete mortality within 24 h after reperfusion, whereas mortality markedly declined to 0% and 10% with shorter (4 h) and longer (6 h) ischaemic durations, respectively. These findings suggest a time-dependent ‘critical window’ of ischaemia that maximally triggers systemic inflammatory responses and fatal crush injury. Blaisdell et al.^24^ further corroborated this observation, showing that systemic release of inflammatory mediators from injured muscle was minimal when the crush period was ≤4 h (owing to limited tissue necrosis) and ≥6 h (attributed to the no-reflow phenomenon caused by irreversible microvascular occlusion). In the clinical setting, when limb ischaemia exceeds 6 h, survival often necessitates amputation of the necrotic extremity. Although the precise ischaemic threshold varies by organ, species, and experimental model, we selected a 5 h crush duration in our rat model to reliably induce severe rhabdomyolysis, systemic inflammation, and lethal CS, consistent with previous reports^5^.

In clinical practice, inhaled NO is commonly administered at concentrations ranging from 10 to 250 ppm for conditions such as idiopathic respiratory distress syndrome (IRDS) in neonates and severe coronavirus disease 2019 in adults^25–27^. In the present study, we evaluated two concentrations of inhaled NO—20 ppm (low) and 160 ppm (high)—to determine their therapeutic efficacy in a rat model of CS. Notably, inhalation of 160 ppm NO significantly improved survival compared with 20 ppm (Fig. 1). Furthermore, administration of 160 ppm NO significantly improved blood pressure parameters (systolic, diastolic, and mean blood pressure), as well as heart rate, particularly when administered after reperfusion, compared with untreated CS control rats (Table 1). These findings suggest that NO-mediated pulmonary vasodilation enhances ventilation–perfusion matching, augments pulmonary venous return, and improves cardiac output and heart rate in the setting of CS-induced circulatory compromise. A recognised clinical concern associated with high-dose NO inhalation is methaemoglobinaemia, which typically warrants discontinuation when Met-Hb levels exceed 5%^28^. In our study, inhalation of 160 ppm NO resulted in peak Met-Hb levels of approximately 4% in the 2 h before and 1 h before /1 h after reperfusion groups and below 3% in the 2 h after reperfusion group (Supplementary Fig. S2), remaining within clinically acceptable limits^29^. The variation in Met-Hb levels appeared to depend on the timing of NO administration relative to reperfusion. During the ischaemic phase, substantial tissue oedema in skeletal muscle likely sequesters erythrocytes and plasma components within interstitial and intracellular spaces, reducing their contribution to systemic circulation. Upon reperfusion, re-entry of these components may dilute circulating Met-Hb concentration. Accordingly, in the before reperfusion groups, Met-Hb levels peaked approximately 1 h after NO exposure and declined rapidly after reperfusion, whereas in the after-reperfusion group, peak Met-Hb levels were attenuated owing to immediate dilution by restored circulation. These observations underscore the safety of 160 ppm NO inhalation with respect to methaemoglobinaemia in this CS model.

Among the NO intervention regimens, inhalation of 160 ppm NO initiated 2 h after reperfusion conferred the greatest survival benefit in rats with CS (Fig. 1b). This timing was associated with the most pronounced improvements in haemodynamics, biochemical parameters, and renal function (Tables 1–3 and Fig. 3c). In CS, the lungs are the first organ exposed to cellular debris and damage-associated molecular patterns released from ischaemic skeletal muscle, triggering interactions with the pulmonary vascular endothelium, neutrophil infiltration, induction of inducible NO synthase, and generation of reactive oxygen species. These processes promote pulmonary inflammation and, in severe cases, ARDS^30,31^. Inhaled NO mitigates this inflammatory cascade by suppressing the expression of adhesion molecules, such as intercellular adhesion molecule-1 on the pulmonary vascular endothelium, thereby limiting neutrophil adhesion and systemic propagation of inflammatory mediators^14^. Furthermore, NO-induced pulmonary vasodilation improves ventilation–perfusion matching, arterial oxygenation, and venous return, ultimately enhancing cardiac output and heart rate in CS rats (Table 1).

We have previously reported that nitrite, an endocrine reservoir of NO under hypoxic conditions, is depleted in ischaemic skeletal muscle of CS rats where it is reduced to NO within mitochondria to support ATP generation under impaired oxidative phosphorylation^12^. This depletion suggests that exogenous NO supplementation—via intravenous administration or inhalation—may be required to protect organs from oxidative stress during reperfusion. In the present study, we focused on inhaled NO as a strategy to deliver NO not only to the pulmonary vasculature but also to distal tissues, including skeletal muscle. Notably, extrapulmonary effects of inhaled NO have been demonstrated in the systemic circulation^32^. Fox-Robichaud et al.^33^ showed that inhaled NO acts at the peripheral microvasculature, exerting anti-adhesive, anti-vasoconstrictive, and anti-permeability effects in NO-depleted tissues. Additional clinical and preclinical studies support systemic benefits of inhaled NO, including attenuation of reperfusion-associated inflammation in ischaemic limbs^32^, reduction of hepatic injury following liver transplantation^34^, and protection against myocardial I/R injury through accumulation of NO metabolites, such as RSNOs in plasma and tissues^17^.

Although NO was previously thought to be rapidly inactivated in pulmonary circulation through oxidation and scavenging by haemoglobin, forming biologically inert nitrate in plasma and erythrocytes^35,36^, accumulating evidence indicates that inhaled NO (the NO· radical) forms stable protein S-nitrosothiols (RSNOs; NO^+^ covalently bound to thiol groups of proteins or low-molecular-weight thiols) in plasma and erythrocytes during pulmonary circulation^37,14^. These RSNOs circulate systemically and mediate cGMP-independent NO signalling via transnitrosylation, thereby conferring anti-inflammatory and cytoprotective effects in remote organs, such as skeletal muscle and kidney^15–17^. The mechanisms underlying the extrapulmonary effects of inhaled NO have been comprehensively reviewed elsewhere^31^.

An additional critical consideration is the timing of NO inhalation relative to reperfusion, because the ability of inhaled NO-derived RSNOs to reach distant ischaemic organs depends on when NO is administered. NO inhalation initiated 2 h before reperfusion failed to effectively deliver RSNOs to erythrocytes and plasma proteins within ischaemic skeletal muscle tissues. In contrast, NO inhalation initiated 2 h after reperfusion allowed RSNOs to be delivered to tissues at the onset of reperfusion, resulting in more robust protective effects. This benefit exceeded that observed with split timing (1 h before /1 h after reperfusion), indicating that NO inhalation initiated immediately after reperfusion onset may represent the optimal strategy to maximise RSNO delivery and improve survival outcomes in CS. The study focused on specific pulmonary and systemic effects of inhaled NO, potentially overlooking other critical factors influencing outcomes in CS, such as the variability in patient responses to treatment and presence of co-morbid conditions. Furthermore, while the portable NO delivery device shows promise for field deployment, its long-term efficacy, safety, and practicality in diverse disaster scenarios have not been clarified. Finally, the integration of inhaled NO with other supportive measures, while theoretically beneficial, necessitates further exploration to establish optimal treatment protocols and timing to maximise clinical outcomes.

## Conclusion

Inhaled NO exerted multifaceted pulmonary and systemic effects through both cGMP-dependent and -independent mechanisms, including reduced pulmonary vascular resistance, decreased thrombotic risk, and suppression of leukocyte–endothelial interactions across vascular beds, ultimately leading to improved survival in CS rats. Importantly, this study highlights the practical advantages of inhaled NO—particularly its simplicity, portability, and non-invasiveness—over conventional intravenous therapies, making it especially attractive for use in disaster scenarios. Although NO inhalation alone was highly effective, further improvements in clinical outcomes may be achievable by integrating this approach with established supportive measures such as aggressive fluid resuscitation and targeted anti-inflammatory therapies.

The portable, controlled NO delivery device employed in this study represents a clinically feasible, field-deployable solution with strong potential to transform early-phase treatment of CS and improve survival in mass-casualty events.

## Methods

### CS animal model

Male Wistar rats (250–300 g) were obtained from Japan SLC (Shizuoka, Japan) and housed under controlled environmental conditions (23 ± 3°C, 55 ± 15% relative humidity) on a 12 h/12 h light–dark cycle, with *ad libitum* access to food and water. Anaesthesia was maintained with inhaled isoflurane (2–5%), and body temperature was maintained using a heating pad throughout the procedure. The CS model was established as previously described by Murata et al.^5^. Briefly, a rubber tourniquet was applied bilaterally to the hind limbs of each rat, wrapped five times around a 2.0-kg metal cylinder, and secured with adhesive tape (Supplementary Fig. S3). After 5 h of compression, the tourniquet was released by cutting the band and removing the device to initiate reperfusion.

### Experimental design

This study consisted of four separate experiments using the CS rat model (Supplementary Fig. S5).

Experiment 1. Survival rates of CS rats inhaling two concentrations of NO (20 and 160 ppm) were assessed at 0, 1, 3, 6, 24, and 48 h after reperfusion. Based on superior survival outcomes with 160 ppm NO, this concentration and three inhalation timing protocols (2 h before, 1 h before /1 h after, and 2 h after reperfusion) were applied in subsequent experiments.

Experiment 2. CS rats inhaled 160 ppm NO for 2 h at different timings relative to reperfusion (2 h before, 1 h before /1 h after, and 2 h after reperfusion). Vital signs, including blood pressure and heart rate, were measured at 3, 24, and 48 h after reperfusion.

Experiment 3. Met-Hb levels were monitored over time following inhalation of 160 ppm NO using the same three timing protocols described above.

Experiment 4. Biochemical parameters were assessed at 3, 24, and 48 h after reperfusion in CS rats treated with 160 ppm NO under the same three timing conditions.

### Device for the controlled release of NO gas

The NO generator used in this study was prepared as described in our previous reports^18,38^. Briefly, two types of nitrite-type layered double hydroxides (NLDHs) were used to deliver low (20 ppm) and high (160 ppm) concentrations of NO to CS rats. For the low-concentration condition (20 ppm), NLDH synthesised via the reconstruction method (RC-NLDH, 400 mg)^38^ was mixed with iron(II) sulphate heptahydrate (FeSO_4_·7H_2_O; 400 mg) using a mortar and pestle. One quarter of the solid mixture (200 mg total) was then loaded into a 6 mL plastic syringe fitted with sponge caps to retain the powder mixture. The NO generator was dried under vacuum overnight and subsequently sealed in a gas-barrier bag (Lamizip® AL-D, Seisannipponsha, Ltd.) together with a zeolite-type strong desiccant (AZ10G-100, As One Corp.). The packaged generator was stored under refrigerated conditions and opened immediately before use. For the high-concentration condition (160 ppm), NLDH synthesised via the anion-exchange method (AE-NLDH; 800 mg)^18^ was mixed with FeSO_4_·7H_2_O (800 mg) using mortar and pestle. One quarter of this solid mixture (400 mg total) was similarly loaded into a 6 mL plastic syringe with sponge caps, dried under vacuum, and packaged in the same manner as described above.

NO generation was initiated by exposure of the NO generator to humid air, which triggered an anion-exchange reaction between interlayer nitrite (NO_2_^-^) and sulphate (SO_4_^2-^), followed by a redox reaction between NO_2_^−^ and ferrous ions (Fe^2+^)^18,38^. The NO generator was connected sequentially to a humidifier (a plastic column containing wet wipes; Wipers S-200, KIMWIPE), an electric pump (GSP-400FT, GASTEC), a nitrogen dioxide (NO_2_) remover (a plastic column containing calcium hydroxide/lithium chloride pellets; Litholyme®, Allied Healthcare Products, Inc.), a syringe filter (pore size, 0.45 μm), and an electrochemical NO sensor (ToxiRAE Pro, RAE Systems), as illustrated in Supplementary Fig. S4. The flow rate of the pump was set to 0.25 L min^−1^ and adjusted as necessary to maintain the desired NO concentration. Notably, the acidic impurity gas NO_2_ is efficiently removed by basic adsorbents such as calcium hydroxide (Ca(OH)_2_), whereas neutral NO passes through unimpeded^39^. NO generation was monitored at room temperature.

### Analysis of haemodynamics, blood gas levels, biochemical parameters, coagulation, and interleukin levels

The HR, SBP, MBP, DBP, and ECG parameters—including QRS interval, PR interval, QT interval, P wave, and T wave—were recorded using a PowerLab data acquisition system (AD Instruments, Nagoya, Japan). The carotid artery was cannulated with a polyethylene catheter (PE-50 tubing) connected to a pressure transducer. Arterial blood samples were collected from each rat via the carotid artery catheter at 3, 6, 24, and 48 h after reperfusion^5^.

Arterial levels of Glu, K^+^, BUN, haematocrit (Hct), pH, partial pressure of oxygen (pO_2_), partial pressure of carbon dioxide (pCO_2_), BE, anion gap (AG), and lactate (Lac) were analysed using an i-STAT 300F blood gas analyser with CG4+ and EC8+ cartridges (FUSO Pharmaceutical Industries, Osaka, Japan). Creatine phosphokinase (CPK) levels were measured using a creatine kinase assay kit (EnzyChrom, BioAssay Systems Co.). Plasma Mb concentrations were determined using a solid-phase enzyme-linked immunosorbent assay (Rat Myoglobin ELISA; Life Diagnostics, West Chester, PA, USA). Met-Hb levels in blood were assessed using a manual spectrophotometric assay based on the disappearance of the Met-Hb absorption peak at 635 nm (pH 6.6), following conversion of Met-Hb to cyan-Met-Hb by neutralised cyanide^40^.

In each experimental group (3, 24, and 48 h after reperfusion; n=6), venous blood and gastrocnemius muscle tissue samples were collected for the measurement of inflammatory cytokines, tissue thiobarbituric acid-reactive substances (TBARS), and MPO activity^41^. Venous blood samples were obtained at 3, 6, 24, and 48 h after reperfusion via a jugular vein catheter^5^. Serum levels of IL-6, IL-10, IL-1β, and TNF-α were measured using Quantikine® ELISA kits (R&D Systems, Inc., Minneapolis, MN, USA). KIM-1 levels were measured using a Rat TIM-1/KIM-1/HAVCR immunoassay (R&D Systems, Inc.), and plasma and urinary creatinine levels were determined using a creatinine colourimetric assay kit (Cayman Chemical, Ann Arbor, MI, USA).

### Histological evaluation

For histological evaluation, tissue samples from skeletal muscle, lung, and kidney were fixed in 10% neutral-buffered formalin, embedded in paraffin, sectioned, and stained with haematoxylin and eosin^5^.

For lung and skeletal muscle tissues, histological variables scored included alveolar and interstitial inflammation, alveolar and interstitial haemorrhage, oedema, atelectasis, and necrosis. Each variable was graded on a 0–4 scale: 0, no injury; 1, injury involving 25% of the field; 2, injury involving 50% of the field; 3, injury involving 75% of the field; and 4, injury involving the entire field. The maximum possible composite score was 28^42^.

Renal injury was scored by calculating the percentage of tubules exhibiting tubular dilation, cast formation, and tubular necrosis, according to a previously described method^43^. For each kidney, 12 cortical tubules from at least four different regions (three tubules per region) were evaluated, with care taken to avoid repeated scoring of different convolutions of the same tubule. Higher scores reflected more severe injury (maximum score per tubule, 7). Points were assigned for tubular epithelial cell flattening (1 point), brush-border loss (1 point), cell membrane bleb formation (1 point), interstitial oedema (1 point), cytoplasmic vacuolisation (1 point), cell necrosis (1 point), and tubular lumen obstruction (1 point).

## Statistical analyses

Data are presented as mean ± standard error of the mean (s.e.m.). Survival rates were analysed using the log-rank test. Differences between groups were assessed using the one-way analysis of variance with Tukey’s honest significant difference test or Tukey’s test. A *p* value <0.05 was considered statistically significant (Statcel 2, 2nd ed. OMS Publishing Inc.).

## Declarations

### Ethical approval

All animal experiments were conducted in accordance with institutional guidelines for animal use and were approved by the Life Science Research Centre of Josai University (approval no. JU18030).

## Acknowledgements

This work was supported by JSPS KAKENHI Grant Number JP22K09148 (to I.M., J.K., and S.I.), a Grant-in-Aid for Scientific Research Presidential Research Grant of Josai University (to I.M. and J.K., 2023–2024), and the World Premier International Research Center Initiative (WPI), MEXT, Japan (to S.I. and N.I.).

## Author contributions

I.M. developed the experimental system for the rat crush syndrome model, performed most of the animal experiments, and generated key data. J.K. proposed the concept of nitric oxide inhalation for crush syndrome and contributed to data interpretation and manuscript preparation together with I.M. S.I. and N.I. developed the nitric oxide gas delivery system and provided it for the experiments. All authors reviewed the manuscript and approved its submission to this journal.

## Supplementary Information

### Supplementary figures

**Supplementary Fig. S1.**
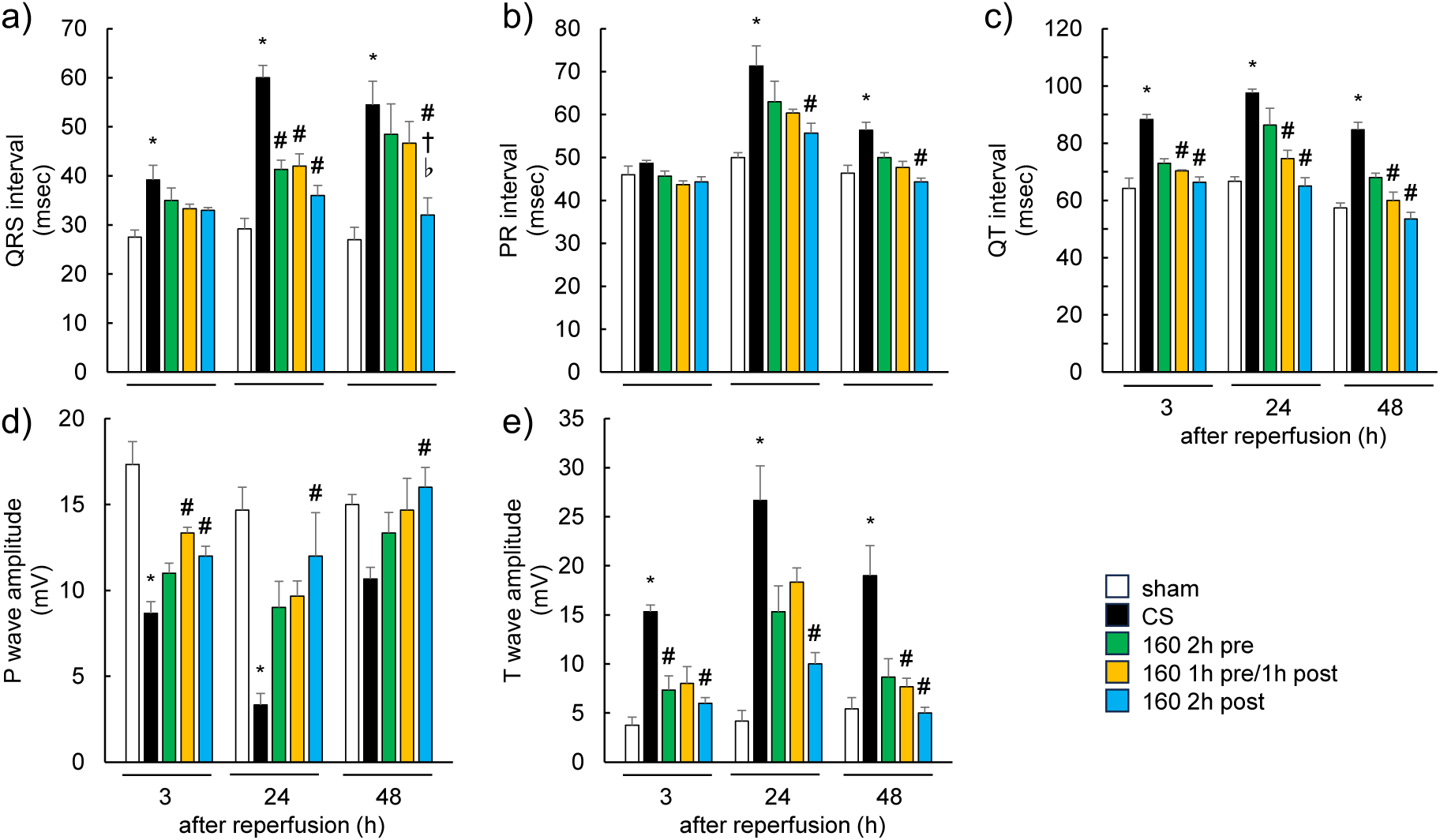
Electrocardiogram parameters of sham, CS control, and NO-inhaled CS rats. ECG parameters measured at 3, 24, and 48 h after reperfusion in sham-operated rats, CS control rats, and CS rats NO for 2 h pre-, 1 h pre-/1 h post-, or 2 h post-reperfusion. The QRS, PR, and QT intervals (Supplementary Figs. S1a–c), as well as T-wave amplitude (Supplementary Fig. S1e), are increased in CS control rats and show a decreasing trend with NO inhalation, particularly in the 2 h post-reperfusion group across all time points. P-wave amplitude is reduced in CS control rats compared with sham rats but increased by NO inhalation, most notably in the 2 h post-reperfusion group (Supplementary Fig. S1d). \**p* < 0.05 vs sham group, #*p* < 0.05 vs CS group, †*p* < 0.05 vs 160 ppm pre-reperfusion group, ♭*p* < 0.05 vs 160 ppm 1 h 160 ppm 1 h pre-/1 h post-reperfusion group (Tukey–Kramer test). Values are presented as the mean ± s.e.m. (n = 6 per group). *Abbreviations*: NO, nitric oxide; ECG, electrocardiogram; CS, crush syndrome; ppl, parts per million; s.e.m., standard error of mean; 160 2 h pre, 160 ppm NO inhalation 2 h before reperfusion; 160 2 h post, 160 ppm NO inhalation 2 h after reperfusion; 160 1 h pre/1 h post, 160 ppm NO inhalation 1 h before and 1 h after reperfusion.

**Supplementary Fig. S2.**
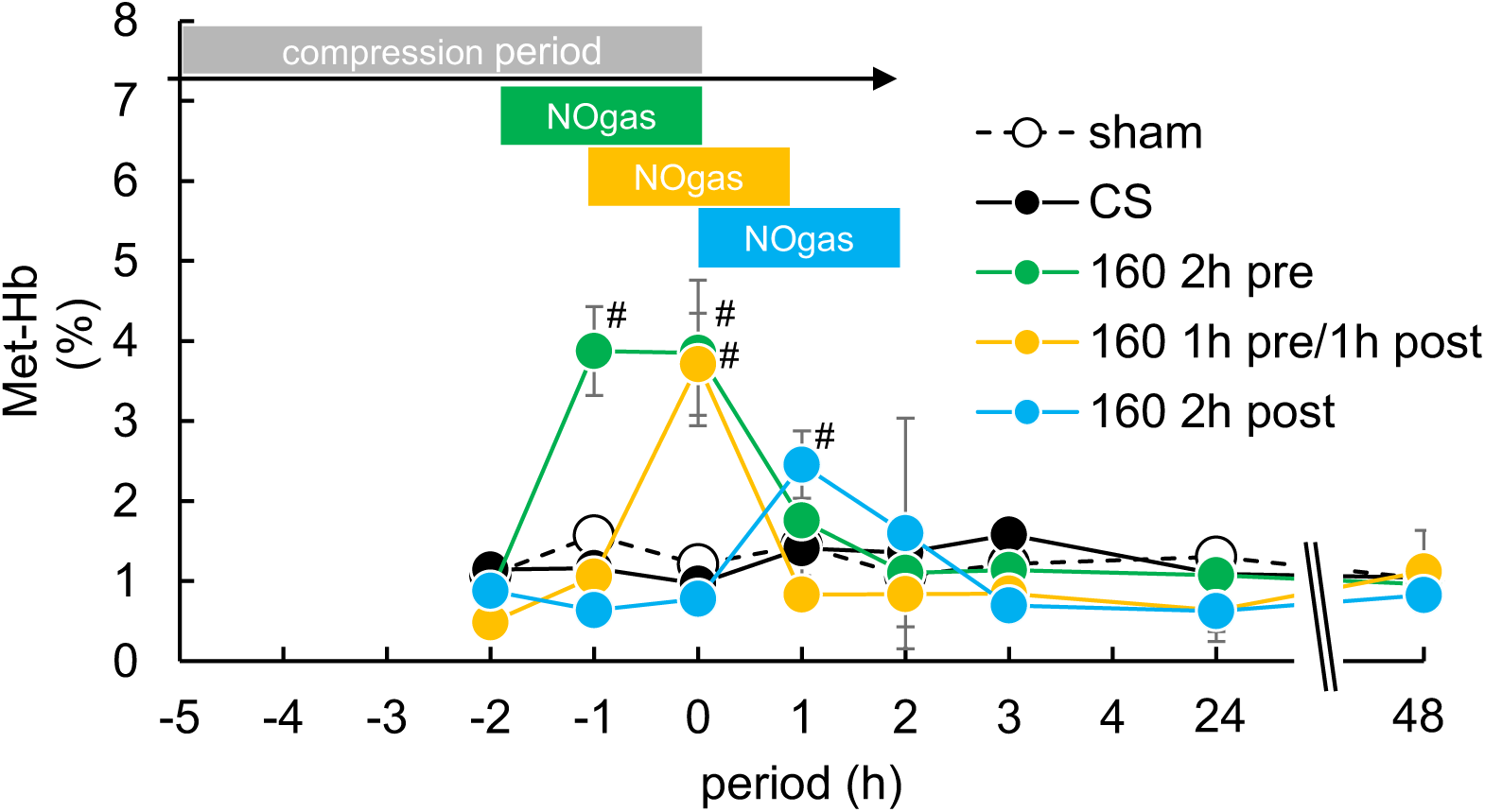
Met-Hb levels of sham, CS controls, and NO inhaled CS rats. The acute phase of Met-Hb levels from 2 h pre-reperfusion to 48 h post-reperfusion show that the inhaled NO at 2 h pre- and 1 h pre-/1 h post-reperfusion reaches approximately 4% of the Met-Hb level. Inhaled NO at 2 h post-reperfusion shows less than 3% Met-Hb from 2 h pre-reperfusion to 48 h post-reperfusion. *Abbreviations*: NO, nitric oxide; CS, crush syndrome; Met-Hb, methaemoglobin; 60 2 h pre, 160 ppm NO inhalation 2 h before reperfusion;160 2 h post, 160 ppm NO inhalation 2 h after reperfusion; 160 1 h pre/1 h post, 160 ppm NO inhalation 1 h before and 1 h after reperfusion.

**Supplementary Fig. S3.**
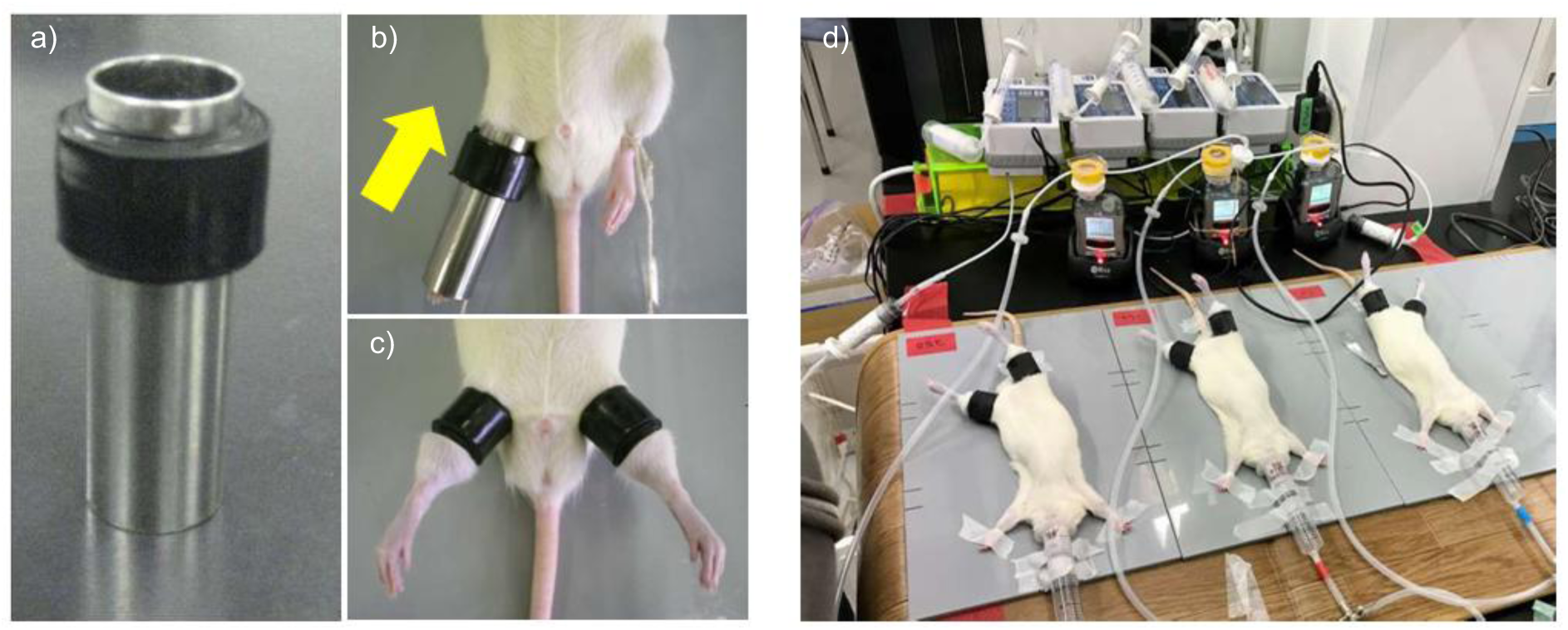
Establishment of the crush syndrome rat model. a) Rubber tourniquet applicator used for hind limb compression. b) Application of tourniquets by sliding in the direction indicated by the arrow. c) Representative positioning of rubber tourniquets on both hind limbs. d) Experimental image showing CS rats receiving inhaled nitric oxide. *Abbreviations*: CS, crush syndrome; NO, nitric oxide.

**Supplementary Fig. S4.**
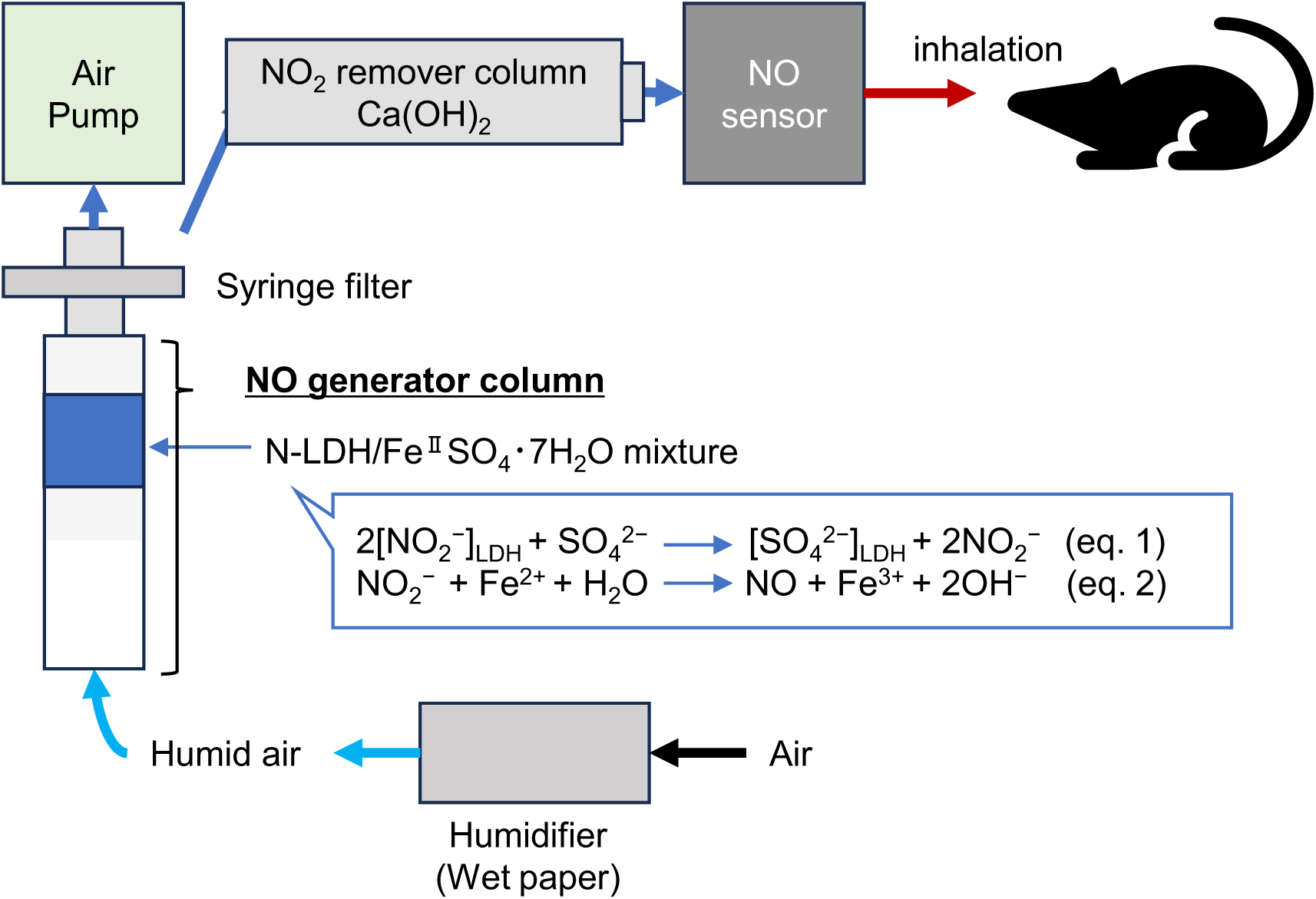
Generation of nitric oxide gas. Schematic representation of the NO generation system. An NO generator (a plastic column containing nitrite-type layered double hydroxides (NLDH)/iron(II) sulphate heptahydrate (FeSO_4_·7H_2_O)) is connected sequentially to a syringe filter (pore size, 0.45 μm), a humidifier (a plastic column containing wet wipes; Wipers S-200, KIMWIPE), an electric pump (GSP-400FT, GASTEC), an NO_2_ remover (a plastic column containing calcium hydroxide (Ca(OH)_2_)), and an NO sensor (ToxiRAE Pro, RAE Systems). The pump flow rate is typically set to 0.25 L/min, and NO generation is monitored at room temperature (20 ± 2 °C). *Abbreviations*: NO, nitric oxide; NO_2_, nitrogen dioxide; NLDH, nitrite-type layered double hydroxide; NO_2_^−^, nitrite ion; FeSO_4_·7H_2_O, iron(II) sulphate heptahydrate; Ca(OH)_2_, calcium hydroxide.

**Supplementary Fig. S5.**
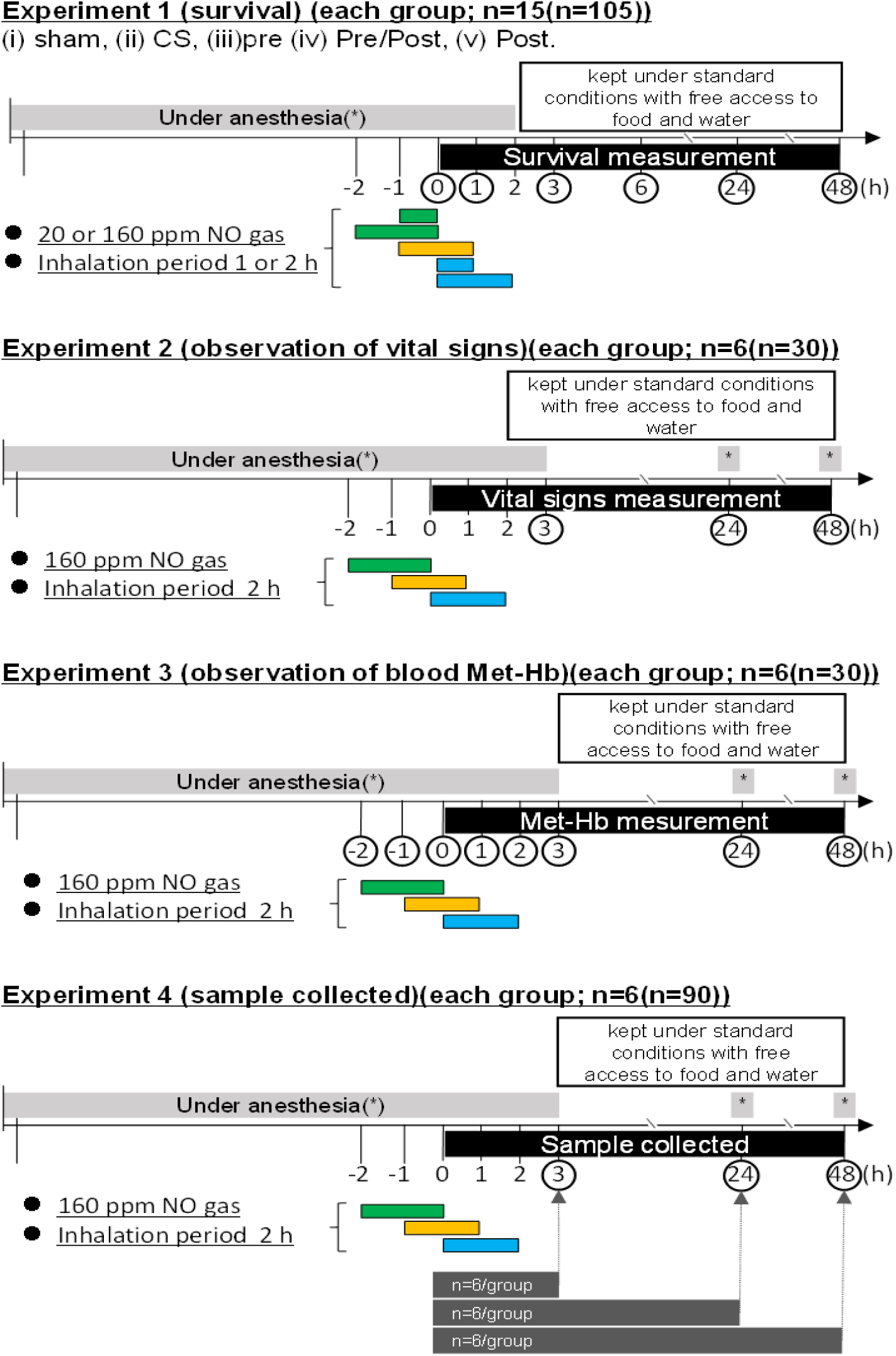
Experimental design of the crush syndrome rat model. Schematic overview of the experimental protocol used in this study, comprising four separate experiments (Experiments 1–4) performed in the CS rat model. *Abbreviations*: CS, crush syndrome; NO, nitric oxide; Met-Hb, methaemoglobin.

